# Identifying Membrane Protein-Lipid Interactions with Lipidomic Lipid Exchange-Mass Spectrometry

**DOI:** 10.1101/2023.06.02.543293

**Authors:** Guozhi Zhang, Melanie T. Odenkirk, Colleen M. Janczak, Ray Lee, Kevin Richardson, Zhihan Wang, Craig A. Aspinwall, Michael T. Marty

## Abstract

Lipids can play important roles in modulating membrane protein structure and function. However, it is challenging to identify natural lipids bound to membrane proteins in complex bilayers. Here, we developed lipidomic lipid exchange-mass spectrometry (LX-MS) to study the lipid affinity for membrane proteins on a lipidomic scale. We first mix membrane protein nanodiscs with empty nanodiscs that have no embedded membrane proteins. After allowing lipids to passively exchange between the two populations, we separate the two types of nanodiscs and perform lipidomic analysis on each with liquid chromatography and mass spectrometry. Enrichment of lipids in the membrane protein nanodiscs reveals the affinity of individual lipids for binding the target membrane protein. We apply this approach to study three membrane proteins. With the *E. coli* ammonium transporter AmtB and aquaporin AqpZ in nanodiscs with *E. coli* polar lipid extract, we detected binding of cardiolipin and phosphatidyl-glycerol lipids to the proteins. With the acetylcholine receptor in nanodiscs with brain polar lipid extract, we discovered a complex set of lipid interactions that depended on the head group and tail composition. Overall, lipidomic LX-MS provides a detailed understanding of the lipid binding affinity and thermodynamics for membrane proteins in complex bilayers and provides a unique perspective on the chemical environment surrounding membrane proteins.

## INTRODUCTION

The lipid composition and organization of membranes differs between tissues, cell types, organelles, and disease states.^1^ Locally, the membrane lipid composition has profound effects on cellular physiology. Lipids can act independently, as signaling molecules and in metabolic pathways.^2^ They can also act collectively, affecting biophysical properties like membrane pressure, fluidity, and thickness.^3^

Through both direct interactions with specific lipids and indirect effects of bilayer properties, lipids can play important roles in regulating membrane proteins.^3-5^ For example, specific binding of phosphatidyl-inositol bisphosphate (PIP_2_) lipids is required for activity of some potassium channels.^6,7^ Conversely, indirect effects of membrane fluidity can influence G-protein coupled receptor activity.^3-5^ These insights have generally been gained by studying membrane proteins in controlled lipid environments, where the effects of specific lipids and bilayer properties can be directly tested. However, it is often unclear how membrane proteins interact with lipids in uncontrolled, heterogeneous, and dynamic natural membranes. In particular, we know that membrane proteins can remodel their surrounding lipids,^8,9^ but we struggle to identify natural lipids bound to membrane proteins in complex bilayer environments.

Here, we present a new approach to identify and quantify the enrichment of natural lipids surrounding membrane proteins in nanodiscs using lipidomic lipid exchange-mass spectrometry (LX-MS). Nanodiscs are nanoscale membrane mimetics that enable membrane proteins to be solubilized in a controllable lipid bilayer.^10,11^ This approach exploits passive diffusion of lipids between nanodiscs to allow lipids to partition between filled nanodiscs that contain a target membrane protein and “empty” nanodiscs that do not (**Figure 1A**).

**Figure 1.**
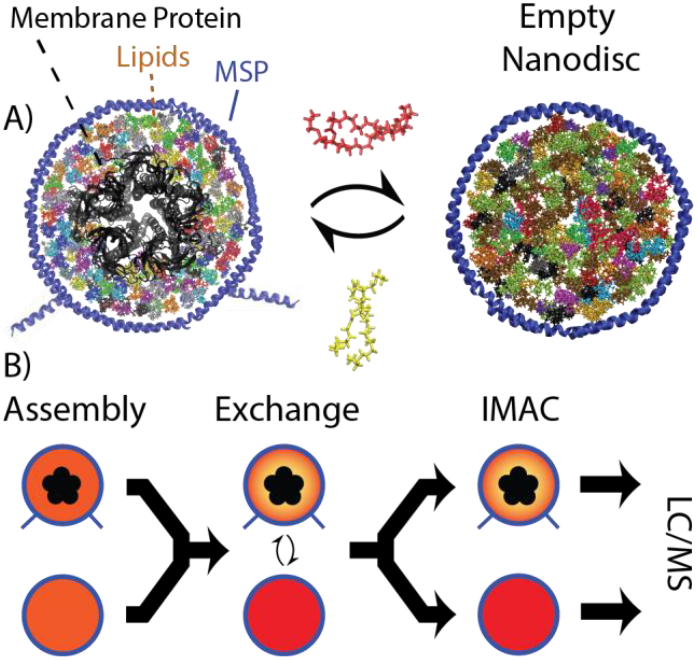
A) Schematic representation of lipid exchange between membrane protein nanodiscs (*left*) and empty nanodiscs (*right*). The membrane protein, AChR, is shown in *black* with the membrane scaffold protein (MSP) in *blue*. Multicolor lipids in (A) indicate the diversity of natural lipids used. B) An overview of the workflow for LX-MS experiments. Mixed *orange* lipids in the assembly are shown conceptually partitioning into *red* empty nanodiscs and a gradient with *yellow* lipids bound to the membrane protein.

Our hypothesis is that lipids that associate with the membrane protein will be enriched in the membrane protein nanodiscs relative to the empty nanodisc. This association is not necessarily a direct protein-lipid contact but rather a preference for the local environment around the protein over the bulk. Previously, we used lipid exchange between nanodiscs to survey the preferential lipid environment of the *E. coli* ammonium channel AmtB with two synthetic lipids.^8^ Barnaba *et al*. used phosphorus NMR to study enrichment of natural lipid classes in membrane protein nanodiscs with peptide scaffolds.^12,13^ Here, we comprehensively characterize bound lipids from a complex natural mixture by developing lipidomic LX-MS (**Figure 1B**). Although nanodiscs do not perfectly replicate the physiological environment, this approach models the complexity of natural lipid mixtures and reveals the affinities of lipid binding on a lipidomic scale and against a complex lipid bilayer background.

To achieve comprehensive lipid head group and tail coverage, mass spectrometry (MS)-based lipidomics provides a powerful tool for identification and quantitation of lipids. However, conventional lipidomics does not capture local interaction information. To address this limitation, several studies have used lipidomics to characterize lipids that co-purify with membrane proteins, either in detergent solutions or within polymer nanoparticles like styrene maleic acid lipid particles (SMALPs).^14-17^ However, detergents may bias lipid binding,^18^ and polymer scaffolds used for nanoparticle formation can affect the lipids that are captured.^19^ In contrast, our recent study indicated that lipoprotein nanodiscs have highly consistent lipidome profiles across different assembly conditions and belt sizes.^20^

Here, we use lipidomics with lipid exchange to characterize how membrane proteins remodel their surrounding environment to enrich specific lipids. We first tested our hypothesis on two bacterial membrane proteins, the *E. coli* AmtB and aquaporin Z (AqpZ), exchanged in *E. coli* polar lipid extract. Both proteins bind cardiolipin (CL) lipids, which are functionally important for AqpZ.^16,21,22^ We then explored the lipid binding preferences of the eukaryotic nicotinic acetylcholine receptor (AChR) in porcine brain polar lipid extracts. With all three systems, we observed distinct lipids enriched around the membrane protein, demonstrating the ability of LX-MS to reveal the lipid environment surrounding membrane proteins and profile lipid binding affinity on a lipidomic scale.

## METHODS

### Membrane Protein Purification

AmtB and AqpZ were expressed in *E. coli* and purified as previously described.^8,23^ Membrane scaffold protein (MSP) MSP1E3D1 (E3) was purified as previously described^11,20^ in either the polyhistidine-tagged (+) or untagged (–) form. Nicotinic acetylcholine receptor (AChR) was purified from *T. californica* electroplaque tissue as previously described.^24^ Experimental details are provided in the Supplemental Methods.

### Nanodiscs Assembly and Purification

AmtB and AqpZ nanodiscs were assembled using E3(−) as previously described.^8,23^ Notably, the MSP used for AmtB and AqpZ nanodiscs had the polyhistidine tag removed. Conversely, AChR nanodiscs were assembled using E3(+) with the polyhistidine tag retained for the scintillation proximity assay.

*E. coli* polar lipid extract (EPL) and brain polar lipid extract (BPL) from Avanti Polar Lipids were dissolved in chloroform, dried under nitrogen gas, and left overnight under vacuum. Dried lipids were quantified with an analytical balance. The average molecular weight of 700 g/mol was used to approximate EPL and BPL concentrations. Weight was used for quantitation rather than phosphate concentration because several classes of brain lipids do not include phosphate groups. Both lipids were dried overnight and resuspended in 100 mM sodium cholate (Sigma-Aldrich) to a final approximate lipid concentration of 50 mM.

Lipids in cholate were then mixed with MSP and AmtB/AqpZ/AChR. Lipids were added at an approximate molar ratio of 120:1 lipid per MSP, using EPL for AmtB/AqpZ and BPL for AChR. AmtB and AqpZ were assembled at a molar ratio of 1 membrane protein complex per 10 nanodiscs. AChR was assembled at a molar ratio of 1:7.5 complex per nanodisc. The mixture was incubated on ice for approximately 1 h before the addition of Amberlite XAD-2 beads (Sigma-Aldrich). The reconstitution mixture was incubated at 4°C overnight on an orbital shaker.

AmtB/AqpZ nanodiscs were purified using immobilized metal affinity chromatography (IMAC) with a HisTrap HP column (Cytiva) to isolate nanodiscs with the embedded membrane protein, which were then further purified by SEC using a Superose 6 10/300 Increase GL (Cytiva). After SEC, nanodiscs were incubated with TEV protease overnight to cleave the HIS-MBP or GFP-HIS tag. Nanodiscs were purified using another round of IMAC to remove the cleaved tag and finally by SEC before being concentrated to 1–3 μM.

For AChR, nanodiscs were purified using only the SEC column because the nanodiscs containing AChR were sufficiently larger than the empty nanodiscs due to the AChR soluble domains. The fractions containing a single AChR complex embedded were collected. After SEC, AChR nanodiscs were concentrated to approximately 1.5 μM for lipid exchange experiments. Unconcentrated nanodiscs were tested with a scintillation proximity assay using nanoSPA™ to confirm activity, as described in the Supplemental Methods.

To test for enrichment of lipids during nanodisc assembly, control EPL E3(−) nanodiscs without membrane protein embedded were obtained during the purification of the AmtB/AqpZ nanodiscs by collecting the flow through of the first IMAC column. Similarly, control BPL E3(+) nanodiscs were isolated by collecting the empty nanodisc peak from the SEC during AChR nanodisc purification. These were reinjected to the SEC and characterized by SDS-PAGE to confirm no membrane protein incorporation.

Donor nanodiscs used in the exchange were assembled as previously described^25^ with either EPL E3(+) or BPL E3(–) and were purified by SEC. The final products of nanodisc assemblies were verified using SDS-PAGE. Nanodiscs assemblies were performed in triplicate, and were aliquoted and stored at –80 °C.

### Lipid Exchange

Lipid exchange was performed at room temperature in 200 mM ammonium acetate at pH 6.8, as previously described.^8^ The lipid exchange duration was extended to 5–7 days to ensure lipids had sufficient time to reach equilibrium. AmtB/AqpZ EPL E3(–) nanodiscs were mixed with empty EPL E3(+) nanodiscs at a 1:1 molar ratio. Prior studies have shown that nanodiscs exchange synthetic lipids over the course of around 2– 3 days,^8^ likely through a monomer diffusion model.^26^ Thus, we do not expect that the membrane protein structure, including the large soluble domains of AChR, significantly affect the exchange. Because AmtB and AqpZ are very stable in nanodiscs,^23^ the mixtures were incubated at room temperature (21–23 °C) for 7 days to ensure the lipid exchange to reach equilibrium. Lateral diffusion within nanodiscs, such as would be observed within physiological membranes, is much faster and is unlikely to contribute to the global kinetics of exchange between nanodiscs. Control experiments were also performed using empty EPL E3(–) nanodiscs mixed with empty EPL E3(+) nanodiscs assembled with the same batch of lipids. For all *E. coli* exchange experiments, the final nanodisc concentrations were 1.3 μM.

For AChR exchange experiments, AChR BPL E3(+) nanodiscs were mixed with empty BPL E3(–) nanodiscs at a 1:1 molar ratio and a final nanodisc concentration of 1.3 μM. The mixtures were incubated at room temperature (21–23 °C) for 5 days to allow the lipid exchange to reach equilibrium. Control experiments were also performed using empty BPL E3(+) mixed with the same batch of empty BPL E3(–) nanodiscs at the same concentration. After exchange, tagged nanodiscs and untagged nanodiscs were separated by IMAC as described in the Supplemental Methods.

### Lipidomics

Prior to LC-MS analysis, samples were dried under vacuum and resuspended in methanol to a lipid concentration of 0.1–1 μM. The EquiSPLASH (Avanti Polar Lipids) internal standard mixture, supplemented with tetra-myristoyl-cardiolipin (TMCL) as a standard for the cardiolipin (CL) class, was added to each sample to an approximately 1 μM final lipid concentration. LC-MS was performed as previously described.^20^ Briefly, approximately 5–10 μL of lipids were injected and separated using a 21-minute trap-and-elute RPLC method using a Waters XBridge C8 Direct Connect HP (10 μm, 2.1×30 mm) followed by separation on a Waters Premier ACQUITY UPLC CSH C18 column (1.7 μm, 1×100 mm). The gradient is provided in **Table S1**. The eluent of these separations was introduced onto a Synapt XS mass spectrometer for acquisition of MS1 mass spectra. MS1 was used for quantitation of lipids species identified using accurate mass and retention time comparisons with a library built using FastDDA and manual curation.^20^ Data was analyzed at both the class and individual lipid levels. Data analysis and statistical procedures are described in the Supporting Information with supplemental data in **Tables S2–S7**. A complete list of identified lipids can be found in **Tables S4** and **S5**.

## RESULTS AND DISCUSSION

Here, we investigated the enrichment of lipids around membrane proteins by allowing natural lipid extracts to exchange and equilibrate between nanodiscs with and without embedded membrane proteins. We first explored two bacterial proteins, AmtB and AqpZ, to understand lipid binding in two prokaryotic systems. Then, we studied the lipid binding of eukaryotic AChR in brain polar lipid extract.

### LX-MS of AmtB and AqpZ in *E. coli* Lipid Nanodiscs

We began by performing control experiments that exchanged *E. coli* polar lipids between two populations that both lacked embedded membrane proteins: empty tagged nanodiscs versus empty untagged nanodiscs. Because the MSP belts were identical except for the presence of the His-tag, which does not interact with the membrane, these nanodiscs should show no difference and serve as a negative control for our experimental design. After exchange and IMAC separation, class-based lipid comparisons showed minor but statistically significant increases in phosphatidyl-glycerol (PG) lipids in the untagged nanodiscs that flowed through the IMAC column and a slight increase in phosphatidyl-ethanolamine (PE) lipids in the tagged nanodiscs that eluted from the column (**Figure 2A** and **Figure S1**). This phenomenon could simply reflect intrinsic variations between samples or measurements. It could also be caused by a slight bias introduced by the IMAC separation after exchange, potentially from the higher imidazole buffer used for elution. Alternatively, it could also indicate unexpected bias in the His-tagged MSP belt. Based on these controls, fold changes of less than 5% were disregarded as insignificant, and statistical significance was then evaluated only for classes with changes that exceeded this cutoff.

**Figure 2.**
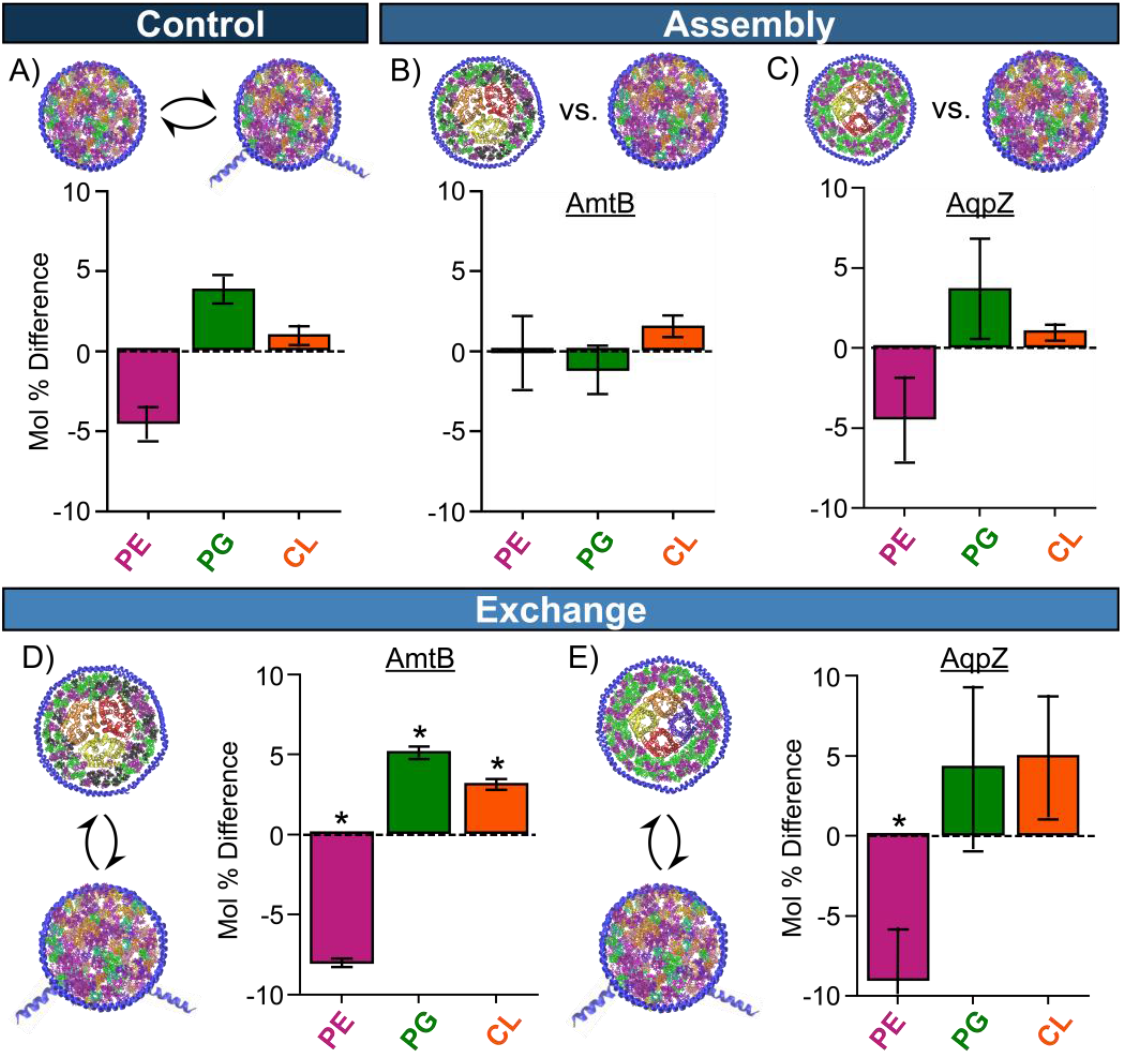
Changes in the *E. coli* lipid class relative molar composition between: (A) empty untagged versus empty tagged nanodiscs without membrane proteins after lipid exchange; nanodiscs assembled with AmtB (B) or AqpZ (C) versus empty untagged nanodiscs prior to lipid exchange; and AmtB (D) or AqpZ (E) untagged nanodiscs versus empty tagged nanodiscs after lipid exchange. Full details are provided in **Table S2**, and absolute mole percentage values are shown in **Figure S1**. Bar heights are the average difference in relative mole percent for each class for three replicates with error bars showing standard deviations. Positive changes represent increases in the empty untagged nanodisc population in A and membrane protein nanodisc population in B–E. Significant (p<0.05) differences that exceed the 5% fold change threshold established by the controls are marked with *. Error bars for PE in (E) that extend off the plot are symmetric

After examining the exchange in empty nanodiscs, we next explored whether AmtB and AqpZ biased the lipids initially incorporated into the nanodisc during assembly, prior to lipid exchange. We considered the lipid class profile changes between nanodiscs with AmtB or AqpZ and empty nanodiscs formed in the same assembly reaction. For both proteins, no significant changes in lipid class were observed prior to exchange (**Figure 2B & 2C**). These results demonstrate that the presence of the AqpZ or AmtB does not bias the *E. coli* lipid profiles during nanodisc assembly.

Finally, we performed LX-MS on both the AmtB and AqpZ nanodiscs. Here, we discovered an enrichment of cardiolipins in both AmtB and AqpZ nanodiscs (**Figure 2D & 2E**). Although there were also significant decreases in PE lipids and increases in PG lipids in the membrane protein nanodiscs, these had similar magnitude to the control and thus may not be due only to the presence of the membrane protein. However, the association of CL with AmtB and AqpZ agrees with prior literature.^22^

We also explored the influence of fatty acid properties on membrane protein preferences, but we observed that within classes, the changes in average fatty acid length and degrees of unsaturation were very small and mostly statistically insignificant (**Figures S2 & S3**). Thus, even though we observed a tendency for CL to exhibit slightly longer carbon chains in AqpZ nanodiscs (+0.07 C for assembly and +0.144 C for exchange), the magnitude of this change was minor.

To explore specific lipid preferences of AmtB and AqpZ, we also measured changes in individual lipids. We first compared the control exchange reaction without membrane proteins to establish a baseline (**Figure S4A**). Using a threshold of p < 0.05 and log_2_ fold change absolute value above 0.5, the control had only 5 out of 52 lipids significantly changing (9%).

For AmtB, 2 lipids (4%) significantly changed during nanodisc assembly (**Figure S4B**), and both were CL lipids. During LX, 10 lipids (19%) changed significantly (**Figure 3A**). Among these species, the greatest fold changes were among several PG lipids (including PG 16:1/16:1, PG 16:1_17:1, and PG 16:1_18:1, although this one also changed in the control) and three CL lipids (CL 72:4, 71:3, and 70:4) that were elevated in AmtB nanodiscs after lipid exchange. In contrast, PE 14:1_16:0 was depleted in AmtB nanodiscs during LX.

**Figure 3.**
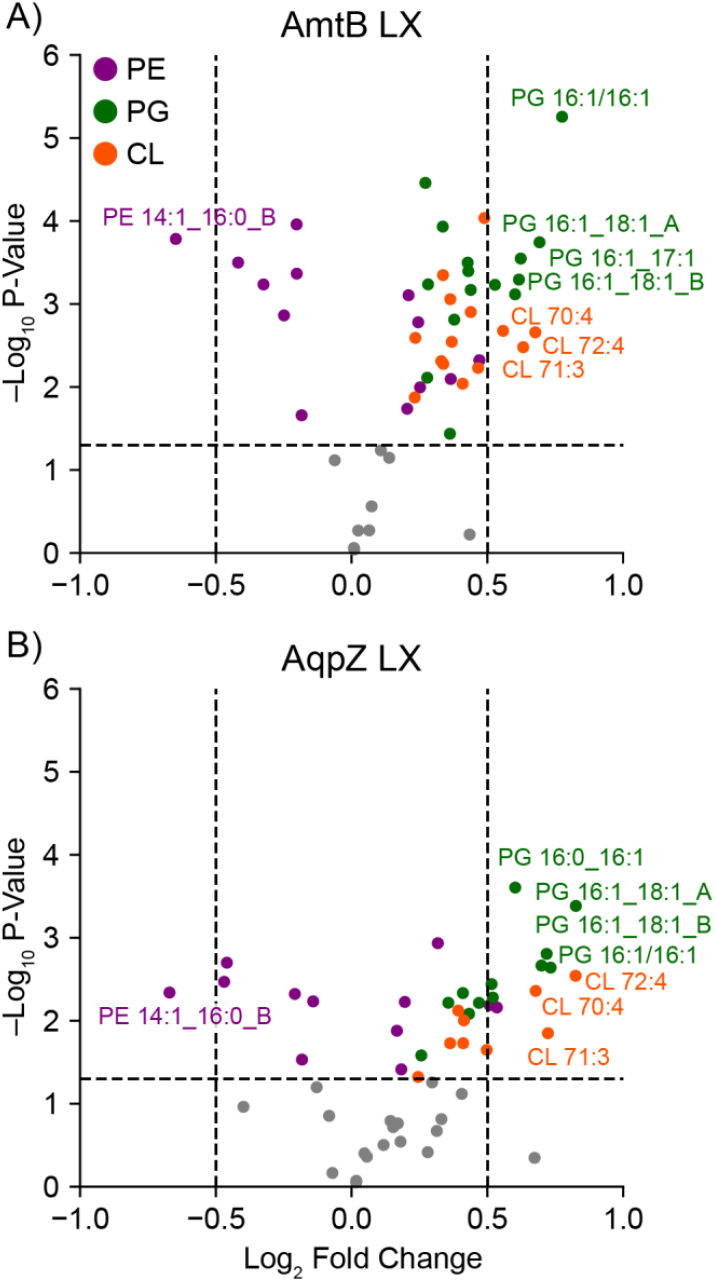
Individual changes in the identified *E. coli* lipids after lipid exchange for AmtB (A) and AqpZ (B). Select lipids with the greatest fold changes and p-values are annotated. Positive log_2_ fold changes indicate higher levels in the membrane protein nanodiscs. Full details are provided in **Table S4**

These results largely agree with the class-based changes showing an enrichment in CL and PG around the protein at the expense of PE lipids. However, they also reveal the specific lipids in each class that interact most favorably with the protein, showing fatty acid details and heterogeneity within classes that were not seen in the class-based analysis.

For AqpZ, no lipids changed significantly during assembly (**Figure S4C**), but 13 lipids (25%) significantly changed during LX (**Figure 3B**). Again, the individual lipid changes during exchange reflected the class-based changes of increased PG and CL lipids in AqpZ nanodiscs and decreased PE lipids but added additional information on fatty acid preferences.

For both proteins, we observed that the bulk of lipid remodeling occurs during the lipid exchange process but not during nanodisc assembly. The tenfold excess of empty nanodiscs during assembly leads to a large pool of lipids that could initially overwhelm any preference of these membrane proteins for specific lipids. However, as discussed below, these data could also reflect that there are different interactions occurring in the detergent environment during assembly than in the bilayer environment during exchange, in this case a weaker overall preference for specific lipids.

Interestingly, the same PG and CL lipid species with the greatest enrichment for AmtB were also among the most enriched in AqpZ. The fact that both AmtB and AqpZ showed preference for the same PGs and CLs suggests these two membrane proteins share a similarity in their lipid binding. Clearly, the anionic head groups of PG and CL are preferred, which is consistent with earlier research, which showed that PG and CL stabilizes their structure and that CL facilitates activity of AqpZ.^22^

It is harder to tell from this data how lipid tails are enriched around each protein. On a class level, the average tail length and unsaturation did not change much between empty nanodiscs and membrane protein nanodiscs during either assembly or exchange. However, individual lipids changes seemed to be most prevalent in species that had 16 or 18 carbon tails. Because these are very common fatty acids, these data suggest that less common shorter or longer fatty acids are not preferred. It may be that both proteins have evolved to prefer the most common lipids in their natural environment.

### Lipid Exchange of AChR in Brain Polar Lipid Nanodiscs

Building on our initial experiments with prokaryotic lipids and membrane proteins, we next extended our lipidomic LX-MS approach to study eukaryotic lipids from brain polar extract, which are much more diverse. Similar to our *E. coli* experiments, we began by testing the control exchange of brain polar lipids across tagged and untagged nanodiscs in the absence of membrane proteins to establish the baseline for class, fatty acid (**Figure 4** and **Figure S5**), and individual lipid (**Figure S6**) perturbations not related to the membrane protein. We assessed the global lipid profiles of the six lipid classes that exhibited sufficient class coverage and reliable internal standard performance: hexosyl-ceramides (HexCer), sphingomyelin (SM), phosphatidyl-choline (PC), phosphatidyl-inositol (PI), phosphatidyl-serine (PS), and PE. There was a small but statistically significant increase in PE and decrease in PS for the eluted His-tagged nanodiscs, but the classes were otherwise identical between the two nanodisc populations (**Figure 4**). The average tail length and unsaturation was also largely unchanged, except for PE lipids. Additionally, the control exchange also had minor but significant differences in the HexCer fatty acid character. With this baseline, we next extended our LX-MS method to probe how adding a membrane protein affected this equilibrium.

**Figure 4.**
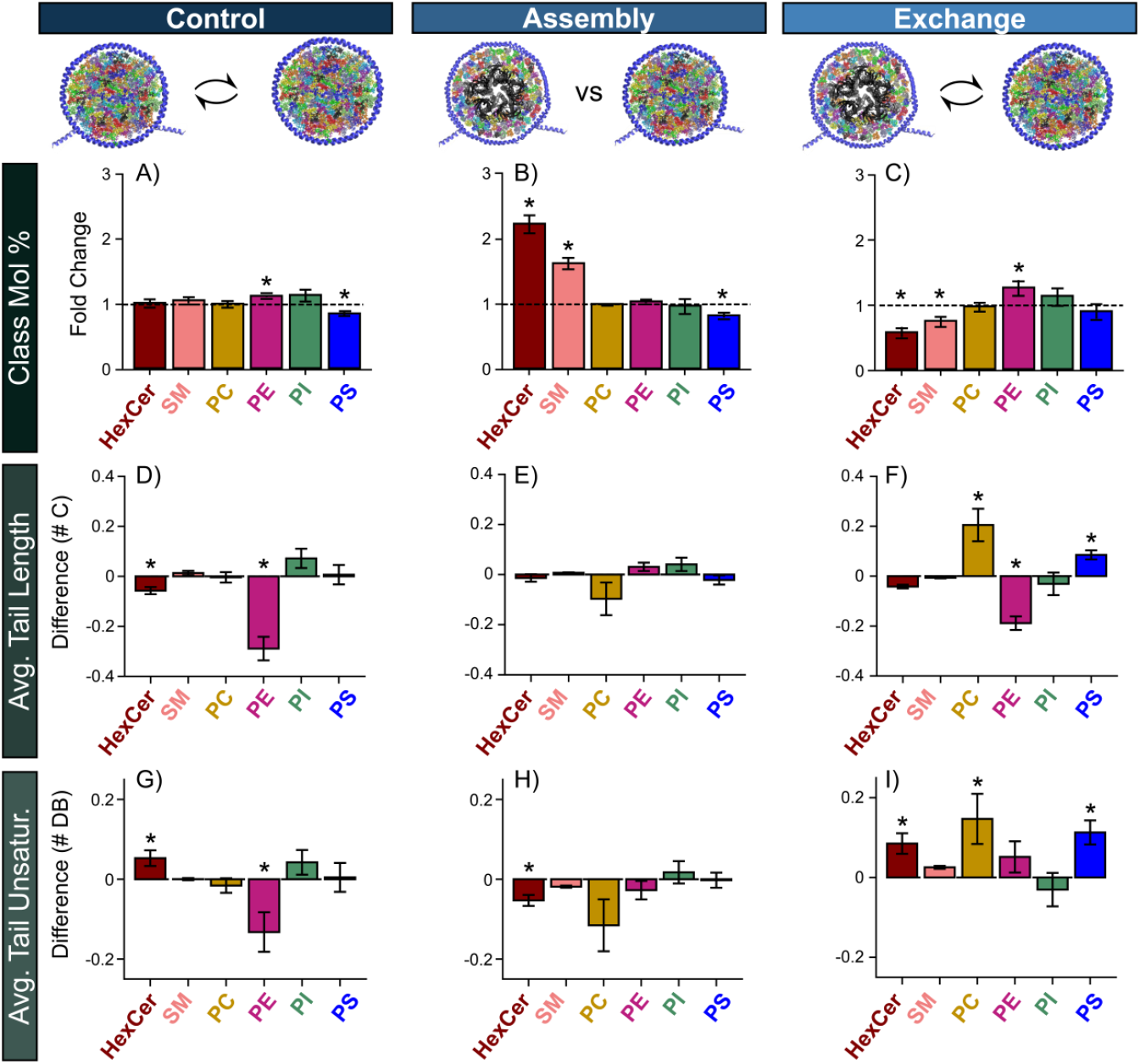
Changes in the brain lipid extract shown as either fold change in relative class mole percent (A–C), difference in the average total number of carbons in the fatty acid tails weighted by the square root intensity (D–F), and difference in the total number of double bonds in the fatty acid tails weighted by the square root intensity (G–I) for: control lipid exchange without AChR (A, D, G), nanodisc assembly (B, E, H), and lipid exchange with AChR (C, F, I). Full results are provided in **Table S3**, and absolute values are shown in **Figure S5**. Bar heights are the average fold change or difference for each class for three replicates with error bars showing standard deviations. Positive changes represent increases in the empty tagged nanodiscs in A, D, and G and AChR nanodiscs in all other panels. Significant (p<0.05) differences that exceed a 5% fold change threshold (A–C) or 0.05 difference (D–I) are marked with *

As our eukaryotic target, we chose the acetylcholine receptor (AChR), which has known functional regulation by the surrounding lipid environment._27_ In experiments with synthetic lipid mixtures, AChR is activated by cholesterol, PE, anionic phospholipids, and sphingolipids._27,28_ However, prior studies have largely focused on synthetic phospholipids. Here, our LX-MS considers AChR and its interaction with brain polar extract lipids to understand protein-lipid interactions in complex eukaryotic lipid environments.

To ensure that AChR was active over the exchange period, we first performed scintillation proximity assays using nanoparticle scintillators (nanoSPA_TM_) to assess the binding of radiolabeled _3_H-acetylcholine._29,30_ AChR in brain lipid nanodiscs bound acetylcholine with a K_d_ of 299 ± 67 nM (**Figure S7**). AChR in POPC nanodiscs had a similar but slightly stronger acetylcholine binding affinity (K_d_=194 ± 53 nM). Both are in line with literature values.^30^ Importantly, we observed no major loss in binding over the course of the lipid exchange time frame (**Figure S8**). Thus, we are confident that AChR remained active during the duration of the lipid exchange.

We first examined differences in lipid class composition in the assembled AChR nanodiscs compared to control nanodiscs to see if AChR biases the lipids incorporated into the nanodisc during assembly, prior to exchange. Unlike AmtB and AqpZ, which showed negligible assembly affects, AChR caused significant changes in lipids during nanodisc assembly. On a class level, the lipid profile of AChR nanodiscs were enriched in HexCer and SM (**Figure 4B**). PS lipids decreased slightly, but this difference was also present within the control.

Interestingly, when we examined the difference in class profiles after lipid exchange (**Figure 4C**), AChR nanodiscs showed a reversal of the assembly effects, with HexCer and SM lipids significantly declining in AChR nanodiscs during assembly. PE lipids were elevated in AChR nanodiscs, but these were also observed in the control.

It is not clear why these sphingolipids would first enrich in AChR nanodiscs during assembly and then flee during exchange. Interestingly, sphingolipids are important for trafficking of AChR,^31,32^ and AChR can cluster in lipid rafts,^33^ which are enriched in HexCer and other sphingolipids.^34^ During nanodisc assembly, lipids and proteins are mixed with detergents (DDM and cholate in this study). It could be that lipids that bind tightly to AChR in detergents are kinetically trapped during nanodisc assembly. At longer equilibration times, lipids could adjust to re-establish a different equilibrium in the context of the new bilayer environment.

Next, we considered how fatty acid length and unsaturation within each class is remodeled by AChR during nanodisc assembly and lipid exchange (**Figure 4D – 4I**). For assembly, no significant trends in fatty acids were observed except for small but significant changes in HexCer unsaturation. However, during lipid exchange, we observed several changes to the fatty acid composition. The class-based average fatty acid length of both PC and PS lipids increased following lipid exchange. Conversely, the average length of PE lipids decreased, but this was also observed in the control. Average class unsaturation also increased slightly with HexCer, PC, and PS lipids. Interestingly, the relative abundance of the PC and PS classes did not change, but their fatty acid composition remodeled to become longer and more unsaturated.

Beyond class-based comparisons, we also considered the individual interactions of the 269 lipids identified with AChR throughout our LX-MS workflow. There were significantly more changes in individual lipids during assembly (**Figure 5A**) and lipid exchange (**Figure 5B**) than observed with controls (**Figure S6**). Specifically, using an absolute log_2_ fold change cutoff of 0.5 and a p-value cutoff of 0.05, only 13 lipids (5%) changed during controls, 57 lipids (21%) changed during assembly, and 119 lipids (44%) changed during lipid exchange. The types of lipids that changed largely agreed with our lipid class observations. For example, during nanodisc assembly, several HexCer lipids were strongly enriched. HexCer lipids then decreased significantly during lipid exchange, mirroring the class-based changes discussed above.

**Figure 5.**
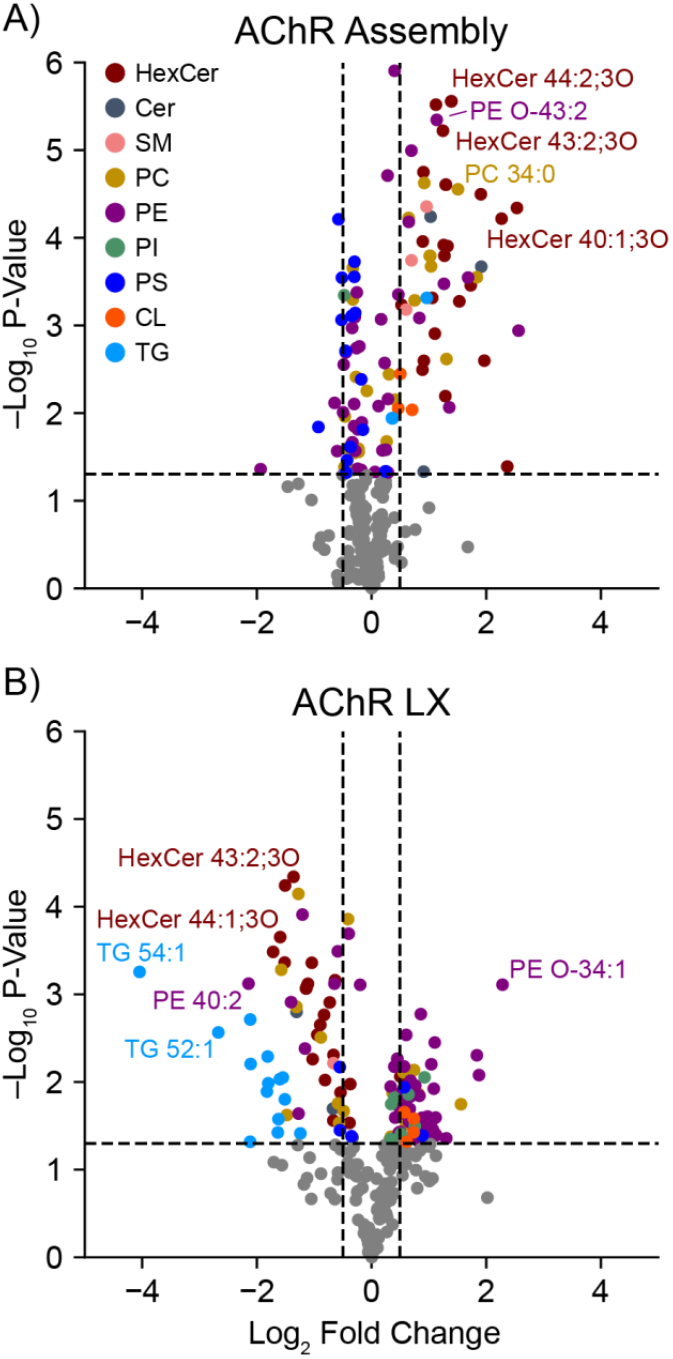
Individual changes in the identified brain lipids during nanodisc assembly (A) and after lipid exchange(B)for AChR. Select lipids with the greatest fold changes and p-values are annotated. Full details are provided in **Table S5**. Positive log_2_ fold change values indicate higher levels in the AChR nanodiscs

Interestingly, triglyceride (TG) lipids did not change during assembly but were significantly decreased during lipid exchange, especially TG 54:1, TG 52:1, and TG 54:5 (**Figure 5**). We hypothesize that TG lipids are kinetically trapped during relatively fast process of nanodisc assembly and are thus initially unchanged. However, given sufficient time to equilibrate, they prefer to be in the empty nanodisc, where they can incorporate into the more flexible hydrophobic interior of the lipid bilayer, like high-density lipoprotein particles that are the basis for MSP nanodiscs.^11,35^

Notably, although anionic lipids did not change significantly as a class, several individual PI and PS lipids changed significantly during assembly and exchange (**Figure 5**), and there were significant fatty acid changes within the PS class (**Figure 4**). Anionic lipids are important for AChR function.^27^ Thus, we infer that anionic PS and PI binding may be more selective, preferring specific tail combinations and binding at specific sites. In line with our observations, binding specific lipids to a small number of sites may not change the overall annular lipid composition dramatically, but it could be reflected in class-based changes to lipid tail composition and/or individual lipids that are enriched around the protein. Unfortunately, both cholesterol and phosphatidic acid (PA) lipids, which are functionally important for AChR,^27^ were not detected by our lipidomic methods, so future studies with alternative MS methods will be needed to explore these. Cholesterol assembles into nanodiscs,^36^ so we expect it is present in these experiments. Overall, our data reveal the dynamic and complex interactions of AChR in bilayers of natural lipids.

### Thermodynamic Interpretation of LX-MS

Until now, we have focused on relative changes in lipid composition between membrane protein nanodiscs and the empty nanodiscs, which provides a qualitative picture of lipid binding preferences to the membrane protein target. However, it is possible to use this data to model the thermodynamics of lipid binding from a few simple assumptions. Biomolecular binding affinity is normally defined with an association constant (P*L ⇌PL), but membrane protein-lipid interactions warrant a slightly different definition because the free protein, P, is always bound to either another lipid or detergent. Thus, affinity can be stated as an exchange of a specific lipid against a generic replacement (PL*L’⇌PL’ *L), as has been previously described.^37-39^ A detailed derivation of binding affinity from the LX-MS data is provided in the Supplemental Methods.

Briefly, we assume that the bulk lipids are fully equilibrated and thus identical in both the membrane protein nanodiscs and the empty nanodiscs. Differences in the membrane protein nanodiscs arise from the annular lipids surrounding the protein. Using known or predicted values for the number of lipids in the annulus versus the bulk lipids in the nanodisc, we can calculate an equilibrium constant for lipid partitioning between the bulk lipids and the annulus, which can be converted into a free energy of lipid binding for each individual lipid (**Figures S9 & S10**). The structure of the free energy plots looks similar but inverted and rescaled from the log_2_ fold change plots (**Figures 3 & 5**).

For AmtB and AqpZ (**Figure S9**), CL and PG lipids that change significantly have negative ΔG values around –2 kJ/mol, and a few PE lipids have positive ΔG in the same range. Free energies are less dramatic for the control exchange and the assemblies. These values are much lower than the –20 to –30 kJ/mol measured for synthetic lipid binding in detergents by native MS,^40^ potentially due to different definitions of affinity, the presence of competing lipids in the bilayer, and/or the complexity of lipid binding in heterogeneous mixtures.^41^ However, they match well with the roughly 2:1 enrichment of PG over PE observed previously with native MS from synthetic lipid nanodiscs, which would translate to around –1.7 kJ/mol.^18^

For AChR, the ΔG values are more extreme, with a cluster of values around –3 to –6 kJ/mol (**Figure S10**). These values are in the same range as previously measured lipid binding energies by EPR for synthetic labeled lipids.^42^ Building on these pioneering studies, our results simultaneously reveal lipid binding affinities for hundreds of individual lipids in a complex natural mixture.

Interestingly, there are a handful of lipids that show infinite or undefined ΔG values. These lipids are so depleted in the membrane protein nanodisc that there would have to be a negative concentration in the predicted lipid annulus to achieve equilibrium of the bulk lipids. Because a negative concentration is not possible, our interpretation is that the membrane protein is affecting the bulk lipids beyond just the annular lipid shell in direct contact with the protein. In other words, the effective lipid annulus is larger for these lipids than previously measured.^42^

For these undefined Δ*G* values and any lipids with positive Δ*G* that are disfavored in the membrane protein nanodiscs, we can calculate the effective annulus size by calculating the size of the annulus at the threshold of producing an undefined value (see Supplemental Methods). Interestingly, the effective annulus for lipids that are significantly depleted in AChR nanodiscs during assembly are clustered around 45, the previously measured size of the annulus (**Figure S11A**).^42^ These data suggest that the influence of the membrane protein on the surrounding lipids does not extend beyond the annulus during nanodisc assembly.

However, after lipid exchange, the effective annulus extends much further for certain types of lipids (**Figure S11B**). For example, there were 22 significant HexCer lipids (76%) that were undefined, which had an effective lipid annulus of 95 ± 31, roughly double the expected size. TG lipids, which strongly prefer the empty nanodiscs for reasons discussed above, had 14 significant lipids (67%) with undefined Δ*G* values, and these had an effective lipid annulus of 145 ± 17, roughly triple the expected size and almost three quarters of the total number of lipids predicted in the nanodisc. Thus, given time to equilibrate, the effective sphere of influence for membrane proteins can extend out into the bilayer beyond the direct lipid contacts and can differ between different lipids.

## CONCLUSION

In summary, we demonstrated the use of nanodiscs with lipidomic LX-MS for comprehensively identifying and quantifying the partitioning of lipids surrounding membrane proteins. This approach enabled the characterization of lipid species with preferential affinity for AmtB, AqpZ, and AChR in bilayers with complex natural lipid mixtures. Profiling the lipid species surrounding membrane proteins will help in understanding the important role that lipids play in regulating membrane proteins. These insights may reveal novel allosteric interactions and uncover essential cofactors required for membrane protein activity.

## ACKNOWLEDGMENT

The authors thank Prof. Stephen Sligar for the MSP plasmids (Addgene #20061) and (Addgene #20066).

## Supporting Information

The Supporting Information contains tables with LC gradient conditions, full results, and statistics available as an .xlsx. The Supporting Information document contains supplemental methods and figures. All raw data for this manuscript is available at MassIVE (DOI: 10.25345/C5599ZB90, MSV000091960).

---

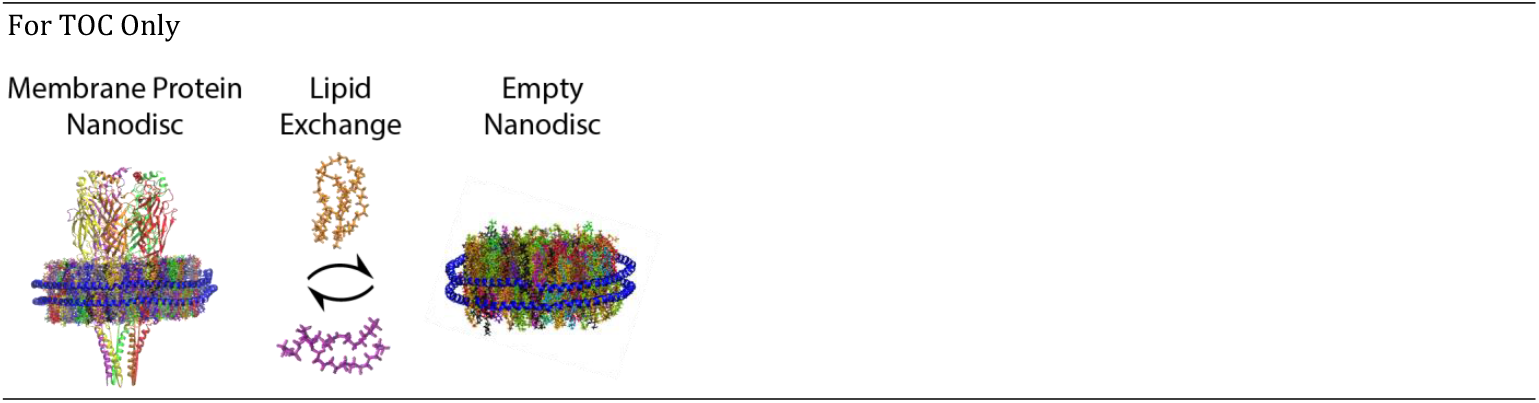

