## Supporting Information for "Identifying Membrane Protein-Lipid Interactions with Lipidomic Lipid Exchange-Mass Spectrometry"

### Supplemental Methods

##### AChR Purification

To purify AChR,[1] approximately 100 g of frozen *T. californica* electroplaque tissue was thawed and homogenized in cold lysis buffer (400 mM NaCl, 20 mM $NaH_2PO_4$ buffer, pH 7.4 with 4 mM N-ethylmaleimide (NEM)) using a blender. Protease inhibitor tablets were dissolved in the homogenized tissue slurry, followed by a centrifugation at 105,000×g at 4 °C for 30 min. The membrane pellets were suspended in 50 mL membrane extraction buffer (80 mM NaCl, 1 mM ethylenediaminetetraacetic acid (EDTA), 20 mM Tris buffer, pH 11.0, with 20% (w/v) sucrose) using a Dounce homogenizer. The homogenized membranes were kept on ice for an additional 30 min to allow for sufficient protein extraction. Membranes were then clarified followed by 3 cycles of centrifugation at 105,000×g for 30 min and resuspended in loading buffer (80 mM NaCl, 20 mM $NaH_2PO_4$, pH 7.4). The pellets were frozen at -80 °C until the next step.

An affinity column for AChR purification was prepared using Affi-Gel 10 (Bio-Rad) that was functionalized with acetylcholine. The beads were prepared by dispersing in 3 different wash buffers followed by mixing on a rotary mixer for 1 h. The first wash buffer, WB1, was 100 mM 4-(2-hydroxyethyl)piperazine-1-ethanesulfonic acid (HEPES) buffer pH 8.0 with 13.3 mM cystamine. WB2 was comprised of 200 mM Tris buffer with 20 mM dithiothreitol (DTT), and WB3 consisted of 200 mM NaCl, 50 mM $NaH_2PO_4$ buffer, pH 7.0 with 20 mM bromo-acetylcholine bromide. The beads were rinsed with 20 mL of water between each wash.

Frozen membrane pellets were thawed on ice in 45 mL loading buffer (80 mM NaCl, 20 mM $NaH_2PO_4$ buffer, pH 7.4). After homogenization, protease inhibitor tablets were added to the membrane suspension, along with the Triton X-100 to a final concentration of 1.5% (v/v). The membrane suspension was mixed on a rotary mixer at 4 °C for 1 h.

The solubilized membranes were then centrifuged at 105,000×g for 30 min. The supernatant solution was diluted with 50 mL of loading buffer, and it was combined with the prepared Affi-Gel 10 beads. Beads and membrane suspension were mixed on a rotary mixer for at least 1 h at 4 °C. Loaded beads were transferred to a 5 mL disposable column. The beads were washed with 50 mL of 80 mM NaCl, 1 mM EDTA, 1 mM *n*-dodecyl β-D-maltoside (DDM), 20 mM Tris buffer, pH 7.4 under gravity flow. AChR was recovered from the column with approximately 30 mL of elution buffer (80 mM NaCl, 1 mM EDTA, 1 mM DDM, 50 mM carbachol, 50 mM βME, 20 mM Tris, pH 7.4). The protein was then buffer exchanged into 20 mM Tris pH 7.4 with 80 mM NaCl and 1 mM EDTA using a 30k MWCO Amicon Ultra-4 centrifugal filter. The final AChR was concentrated to approximately 17.6 nmol in 1.5 mL for nanodisc preparation.

##### Post-Exchange IMAC Separation.

After exchanging, tagged nanodiscs and untagged nanodiscs were separated by immobilized metal affinity chromatography (IMAC) using a Ni-NTA His SpinTrap (GE Healthcare).[2] To ensure the same column volumes for all separations, 350 µL of the Ni-NTA bead suspension were loaded on to the trap. After removal of the storage 20% ethanol, the column volume for each spin trap was estimated to be 175 µL. The SpinTrap was prepared by performing three washes of 600 µL nickel loading buffer (5 mM imidazole, 200 mM ammonium acetate, pH 7.6) to equilibrate the column. To limit non-specific binding, the imidazole concentration of the exchange mixture was adjusted to 5 mM by diluting the exchange mixture with a 10 mM imidazole solution (pH 7.6 adjusted with acetic acid) to limit the non-specific binding. For AChR, this step was not completed due to the low starting concentration of lipids in these samples.

For the separation of lipid exchange nanodisc populations, the total 30 µL of AmtB and AqpZ exchange reaction mixture was added to the IMAC beads and allowed to incubate for approximately 1 min at 4 °C. The flow-through (untagged nanodiscs) were removed by centrifugation at 100×g for 45 seconds. Extra nickel loading buffer (100 μL) was added to the column and washed 3 more times. The collected flow-through and washes contained the non-tagged nanodiscs. The elution of tagged nanodiscs were also collected by adding 50 μL of the nickel elution buffer (500 mM imidazole, 200 mM ammonium acetate, pH 7) followed by 3 washes with 150 μL of nickel elution buffer. After separation, the His SpinTrap was regenerated by washing 3 times with 600 μL aliquots of loading buffer. For AChR, the same steps were followed only with 15 μL of the undiluted exchange reaction mixture due to the low lipid concentrations.

**Scintillation Assay.**

Scintillation proximity assays were completed using nanoSPA™ (Scintillation Nanotechnologies, Inc.).[3] SEC-purified nanodiscs in 200 mM ammonium acetate buffer were combined with 4–10 mg NTA-functionalized nanoSPA™ for His-tag binding. The nanoSPA™ was rinsed 3 times with 10 mM HEPES by centrifuging the particles at 4,700×g for 10 min, removing the supernatant buffer, and dispersing the nanoSPA™ in fresh buffer. After the third wash, the supernatant buffer was removed and replaced with nanodiscs in 200 mM ammonium acetate. The nanoSPA™ with nanodiscs were then gently mixed on a rotary mixer at 4 °C overnight.

Prior to the assay, nanodisc-loaded nanoSPA™ beads were diluted to 0.4–1 mg/mL with assay buffer (125 mM NaCl, 2.5 mM KCl, 1.5 mM $CaCl_2$, 2 mM $MgCl_2$, 2.5 mM phosphate, and 10 mM HEPES, pH 7.4). 3 mL of each sample was placed in 7 mL polyethylene scintillation vials. $^3$H-acetylcholine iodide was combined with unlabeled acetylcholine bromide to prepare a stock concentration of 20 nCi/μL, 10 μM total acetylcholine. When assays were performed in microplates, this stock solution was prepared at 2 nCi/μL, 1 μM total acetylcholine. Small volumes (5–25 μL) of the stock acetylcholine solution were sequentially added to nanodisc-loaded nanoSPA™ samples, allowed to bind for at least 3 min with periodic, gentle mixing, and then counted. Scintillation counts for all proximity assays were measured using a Beckman LS 6000IC scintillation counter set to a wide window (0–2000 keV) for 1-min count times. When assays were performed in microplates, stock acetylcholine was added to the wells to prepare wells with acetylcholine concentrations from 0 to 517 nM. Assay buffer was added to wells when needed to make the final volume of each well 200 μL. Samples were mixed within the plate wells and scintillation counts were measured using a MicroBeta 2 (Perkin Elmer) scintillation counter, again for one-min count times. The timed decay for each sample at the highest acetylcholine concentration was measured to ensure the protein was functional during the duration of the exchange.

**Lipidomics Data Analysis.**

Previously, *E. coli* and brain polar lipid extracts were analyzed with a FastDDA method under the same instrumentation conditions to curate lipid libraries of 55 and 279 unique species.[4] These same libraries were employed here to facilitate accurate mass and time matching of lipids from MS1 acquisitions in Skyline.[5,6] The individual precursor lipid areas were defined for each lipid and exported. Each experiment was performed on three replicate nanodisc assemblies.

Prior to statistical evaluation, data was first assessed for outliers with a Pearson correlation, principal component analysis, and robust Mahalanobis distance test.[7] Any samples that failed to pass all three tests were removed. Additionally, any lipid identifications that did not meet criteria for reliable quantitation were also removed. This resulted in a slight downsizing of lipid identifications to include 52 lipids in *E. coli* and 269 in brain. Following outlier assessment, data was processed by first normalizing to the summed ion current of each sample (mTIC) and transforming the data into $\log_2$ space to generate the volcano plots for individual lipid analysis.

For these statistical comparisons of individual lipid differences, an unpaired ANOVA analysis at a type one error rate of 95% was used to assess relative lipid area fluctuations across each comparison. An additional Holm correction was applied to adjust significance findings for multiple comparison.[7] Results from these comparisons are presented in Supplemental **Table S4** & **S5**.

To elucidate membrane protein-lipid preferences, the data was evaluated across lipid classes by assessing global class changes and fluctuations in fatty acid character (tail length and unsaturation) within each lipid class. Changes in relative lipid class abundances were determined by summing the total class intensity in each class (in normal space) and normalizing it to the internal standard. The total normalized intensity for each class was then divided by the total normalized intensity for all classes. A few lipid classes were excluded due to high variability (LPC and LPE), having only a very small number of lipids (PG, Cer, CL), or for having unreliable internal standard signal (TG). Plasmalogen and alkyl ether lipids were grouped with their respective head group classes. The average lipid tail length and lipid tail unsaturation for each class was determined by taking the average number of carbons on the tail or average number of double bonds weighted by the square root of the signal intensities. A square root weighting was used to avoid having the class too heavily influenced by a few highly abundant lipids.

After determining the relative class abundance, tail length, and tail unsaturation for each sample, three select comparisons were made. First, the control nanodiscs without embedded membrane proteins were compared after lipid exchange between the tagged nanodiscs captured/eluted from the Ni-NTA His SpinTrap and the untagged nanodiscs that flowed through the column. This control comparison reveals any potential measurement or exchange biases between the flow-through (untagged) and eluted (tagged) nanodiscs in the absence of membrane proteins.

Second, the assembled membrane protein nanodiscs prior to mixing or exchange were compared with empty nanodiscs without the protein. These empty nanodiscs were collected from the same assembly reaction by capturing the nanodiscs without the protein that flowed through either the IMAC (AmtB and AqpZ) or the SEC (AChR) column after assembly. This comparison measured any differences in lipid profiles created during assembly by incorporating the membrane protein.

Finally, we compared the tagged and untagged nanodiscs after lipid exchange and separation by IMAC. AChR was assembled with tagged nanodiscs and was thus present in the elution. AmtB and AqpZ were assembled with untagged nanodiscs and were in the flow through. This comparison measures the enrichment of lipids in the membrane protein nanodiscs after lipid exchange.

For each comparison, the fold change was calculated for each lipid class relative abundance, and the absolute differences were calculated for each class weighted average tail length and tail unsaturation. Any fold changes less than 5% were ignored, and any absolute difference in tail length/unsaturation less than 0.05 were ignored. For changes of sufficient magnitude, p-values were calculated by t-tests and were corrected by a Benjamini-Hochberg procedure. Outputs from the global lipid structural changes are presented in **Table S2** & **S3.**

##### Thermodynamics Calculations

We begin by considering the exchange of a specific lipid species, L', against all other possible lipids, L. After lipid exchange between two nanodisc populations, 1 and 2, we can write this as an equilibrium reaction:

$$L'_1 + L_2 \rightleftharpoons L'_2 + L_1 \qquad \textit{Eq. S1}$$

For empty nanodiscs, the equilibrium constant for this reaction is defined as:

$$K = \frac{[L'_2][L_1]}{[L_2][L'_1]} \qquad \textit{Eq. S2}$$

Within this equation, $[L'_2]/[L_2]$ is nearly equal to the mole percent for L' in nanodisc 2, $X'_2$, and can be approximated as the mTIC normalized intensity for L' in nanodisc 2. Similarly, $[L'_1]/[L_1]$ can be expressed as the mole percent of L' in nanodisc 1, $X'_1$, and can be approximated as the normalized intensity for L' in nanodisc 1. Any biases in ionization efficiency should apply equally to both ratios and cancel out. Substituting these values into *Eq. S2*, K can be approximated as the fold change in normalized intensity for L' in nanodisc 2 compared to nanodisc 1, $K = X'_2/X'_1$.

After calculating K, the free energy of exchange can then be calculated as:

$$\Delta G = -RT \ln K \qquad Eq.\ S3$$

where T = 22 °C or 295.15 K, and R is 8.314 × $10^{-3}$ kJ/molK to give units of kJ/mol. For exchange of control nanodiscs with no membrane proteins, ΔG values were calculated for all lipids by substituting their fold change values into *Eq. S3* as K.

For membrane protein nanodiscs, a similar equilibrium constant could be calculated for the exchange of lipids between the two populations of nanodiscs. However, there is a second equilibrium occurring inside the membrane protein nanodiscs between the lipids in the bulk, B, and lipids in the annular belt surrounding the membrane protein, A:

$$L'_B + L_A \rightleftharpoons L'_A + L_B \qquad Eq.\ S4$$

This reaction provides a simplified expression for membrane protein affinity of lipid binding.[8] More complex models have been developed for multivalent exchange where binding one larger lipid may displace multiple smaller lipids, which may be more appropriate for larger lipids like cardiolipin that likely displace multiple smaller phospholipids.[9] For simplicity, we have not applied these corrections here and assume equal valency in exchange.

Using the same derivation as above, we can express the equilibrium constant for this reaction as a ratio of the mole percent of L' in the annulus, $X'_A$, versus the bulk, $X'_B$.

$$K = X'_A / X'_B \qquad Eq.\ S5$$

Although we cannot measure $X'_A$ or $X'_B$ directly, we can calculate them from $X'_1$ and $X'_2$ using a few assumptions. For clarity, we will specify nanodisc 1 as empty, $X'_E$, and nanodisc 2 as the nanodisc population with the membrane protein, $X'_P$. Our key assumption is that the bulk lipids are equilibrated and identical in composition between the two nanodiscs. Because the empty nanodisc is entirely bulk lipids, we can thus conclude that:

$$X'_B = X'_E \qquad Eq.\ S6$$

The membrane protein nanodisc contains a mixture of bulk and annular lipids. We will define *p* as the fraction of bulk lipids in the nanodisc such that:

$$X'_P = p\, X'_B + (1-p)\, X'_A \qquad Eq.\ S7$$

Substituting in *Eq. S6* and rearranging *Eq. S7* to solve for $X'_A$ yields:

$$X'_A = X'_P \frac{1}{1-p} - X'_E \frac{p}{1-p} \qquad Eq.\ S8$$

Substituting *Eq. S8* and *S6* back into *S5* yields:

$$K = \left(X'_P \frac{1}{1-p} - X'_E \frac{p}{1-p}\right) / X'_E = \frac{X'_P}{X'_E}\frac{1}{1-p} - \frac{p}{1-p} \qquad Eq.\ S9$$

As above, we can closely approximate $X'_P / X'_E$ as the fold change in normalized signal intensity for the membrane protein nanodiscs compared to the empty nanodiscs, which is provided from the data. Thus, the only remaining term to specify is *p*, the ratio of bulk lipids in the membrane protein nanodisc.

To calculate *p*, we can rely on published or simulated values for both the total number of lipids in the membrane protein nanodisc and the total number of lipids in the annulus surrounding the membrane protein. For AmtB and AqpZ, the total number of lipids in the nanodisc has previously been determined with native MS to be around 150.[10] Similarly, native MS and molecular dynamics simulations show the lipid annulus to be around 75 lipids for each protein.[11] Thus, the fraction of bulk lipids in both AmtB and AqpZ nanodiscs is 75/150=0.5.

AChR is slightly smaller and thus has both more lipids in the nanodisc and fewer lipids in the annulus. Using CHARMM-GUI[12] to assemble a model nanodisc with AChR (PDB ID: 2BG9)[13], we estimated around 200 lipids in the nanodisc. The number of annular lipids for AChR has previously been measured by EPR at around 45 lipids.[14] Thus, the fraction of bulk lipids in AChR nanodiscs is 155/200=0.775.

Substituting the measured fold changes with the calculated bulk lipid fractions into *Eq. S9* allowed calculation of the equilibrium constant, *Eq. S5*. Adding this with the experimental temperature in *Eq. S3* allows us to calculate the free energy change for L' binding to the membrane protein for each lipid identified in the lipidomics experiment. Results are provided in **Tables S6** & **S7**.

If accurate mole percentages of $X'_E$ and $X'_P$ are measured, these can be substituted with $p$ into *Eq. S8* to directly determine the composition of the lipid annulus. However, these calculations rely on first quantifying the molar ratios within each sample set, which are harder to measure accurately than fold changes across samples sets, due to potential biases in ionization efficiency. Future work will explore these calculations in more detail.

In several cases, specific lipids yielded negative K values in *Eq. S9*, which would yield undefined or infinite $\Delta G$ values in *Eq. S3*. These lipids clearly do not bind the membrane protein, and our interpretation is that the undefined values indicate that the effective lipid annulus is larger for these lipids than previously reported. In other words, the sphere of influence of the protein extends beyond the reported lipid annulus in direct contact with the protein.

To calculate the effective annulus for any lipid that is disfavored for protein binding (K < 1), we can consider the limit where K approaches 0 for a given lipid species. At K=0, the lipid is entirely excluded from the effective annulus, and *Eq. S9* becomes:

$$0 = \frac{X'_P}{X'_E}\frac{1}{1-p} - \frac{p}{1-p}$$

which simplifies to:

$$\frac{X'_P}{X'_E} = p \qquad \textit{Eq. S10}$$

Here, $p$ is defined as the fraction of bulk lipids in the nanodisc, which is $(n_D - n_A)/n_D$, where $n_D$ is the total number of lipids in the nanodisc and $n_A$ is the number of annular lipids around the protein. After calculating $p$ from *Eq. S10*, $n_A$ can be determined with: $n_A = (1 - p)\, n_D$, using the predicted total number of lipids in the nanodisc described above. Individual effective lipid annulus sizes were calculated for each lipid with K < 1. Means and standard deviations were determined for lipids in each class with significant p-values and undefined $\Delta G$ values.

For lipids with K=1, the size of the effective annular lipids is 0, indicating that these lipids do not have any preference for the environment surrounding the membrane protein versus the bulk. The protein has no effect on these lipids. For lipids that are enriched in the lipid annulus (K > 1), these calculations yield effective annulus sizes with negative values, and it is unclear if these values have any physical interpretation. We have chosen to exclude these from calculations of effective annulus.

### Supplemental Figures

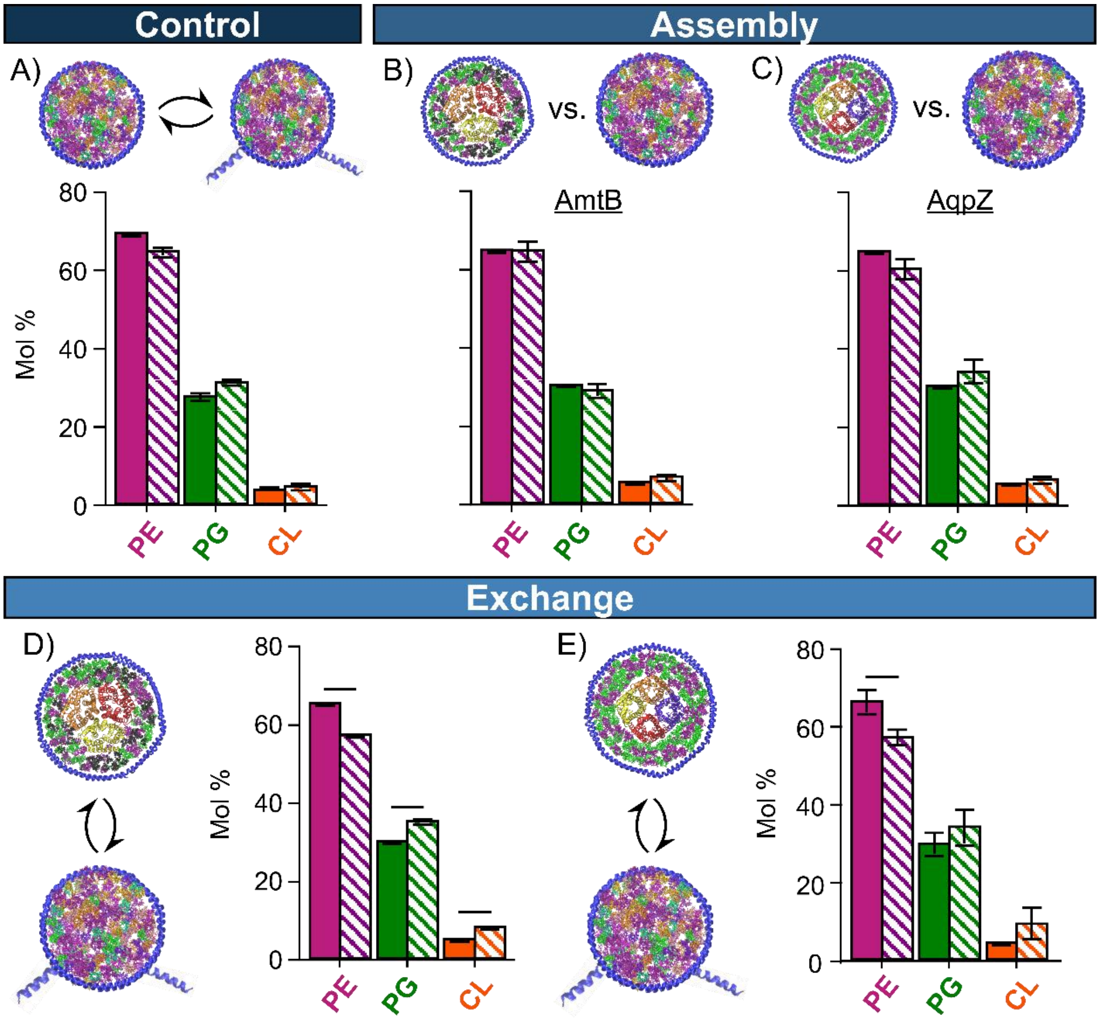

**Figure S1**. Changes in the E. coli lipid class mole percent for: (A) empty untagged (*shaded*) versus empty tagged nanodiscs (*solid*) without membrane proteins after lipid exchange; nanodiscs assembled with (B) AmtB or (C) AqpZ (*shaded*) versus empty untagged nanodiscs (*solid*) prior to lipid exchange; and AmtB (D) or AqpZ (E) untagged nanodiscs (*shaded*) versus empty tagged nanodiscs (*solid*) after lipid exchange. Full details are provided in **Table S2**. Bar heights are the average mole percent for three replicates with error bars showing standard deviations. Significant ($p<0.05$) differences that exceed a 5% fold change threshold are marked with horizontal bars.

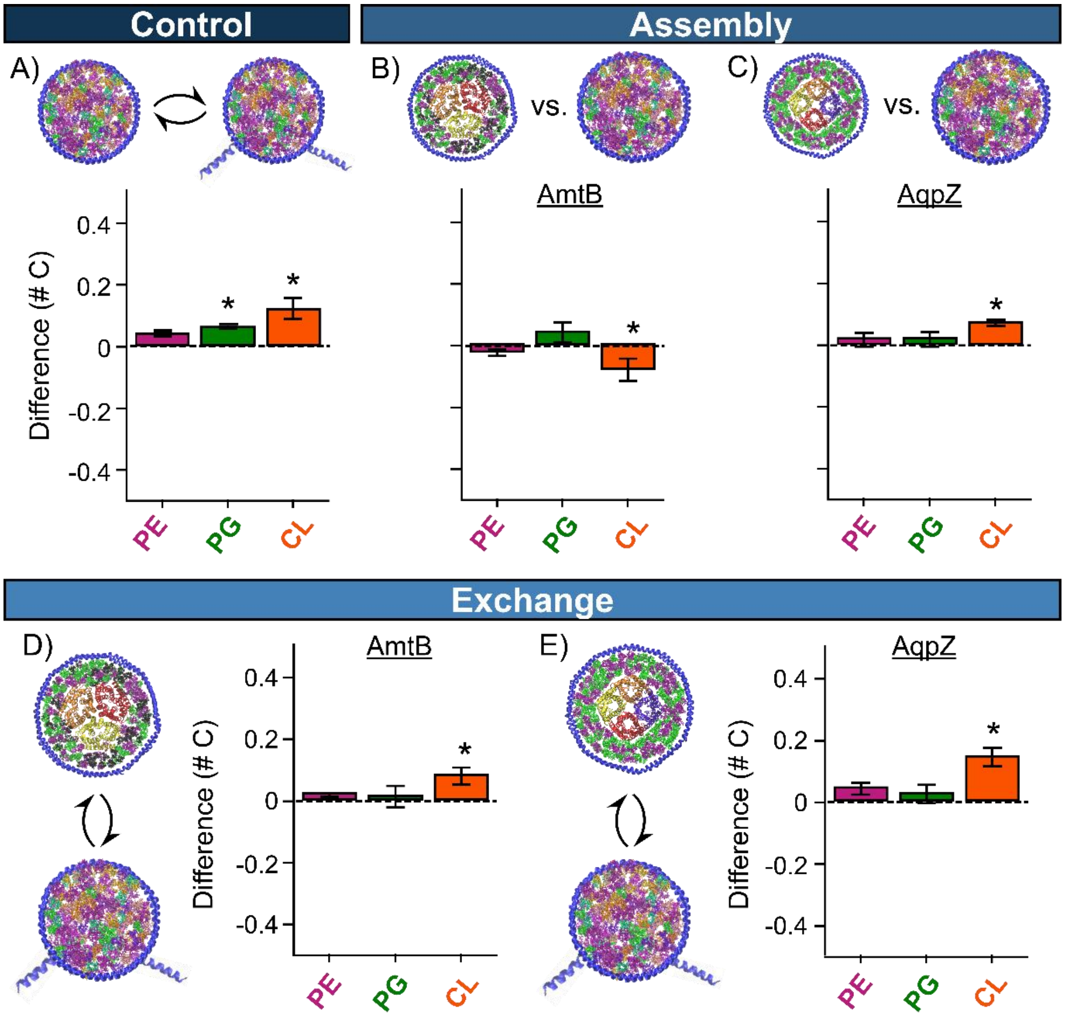

**Figure S2**. Changes in the *E. coli* lipid class average lipid tail length for each class weighted by the square root of their intensity for: (A) empty untagged versus empty tagged nanodiscs without membrane proteins after lipid exchange; nanodiscs assembled with AmtB (B) or AqpZ (C) versus empty untagged nanodiscs prior to lipid exchange; and AmtB (D) or AqpZ (E) untagged nanodiscs versus empty tagged nanodiscs after lipid exchange. Full details are provided in **Table S2**. Bar heights are the average difference in weighted average tail length expressed as total number of carbons for each class for three replicates with error bars showing standard deviations. Positive changes represent longer tails in the empty untagged nanodisc population in A and membrane protein nanodisc population in B–E. Significant ($p<0.05$) differences that exceed a 0.05 carbon difference threshold are marked with *.

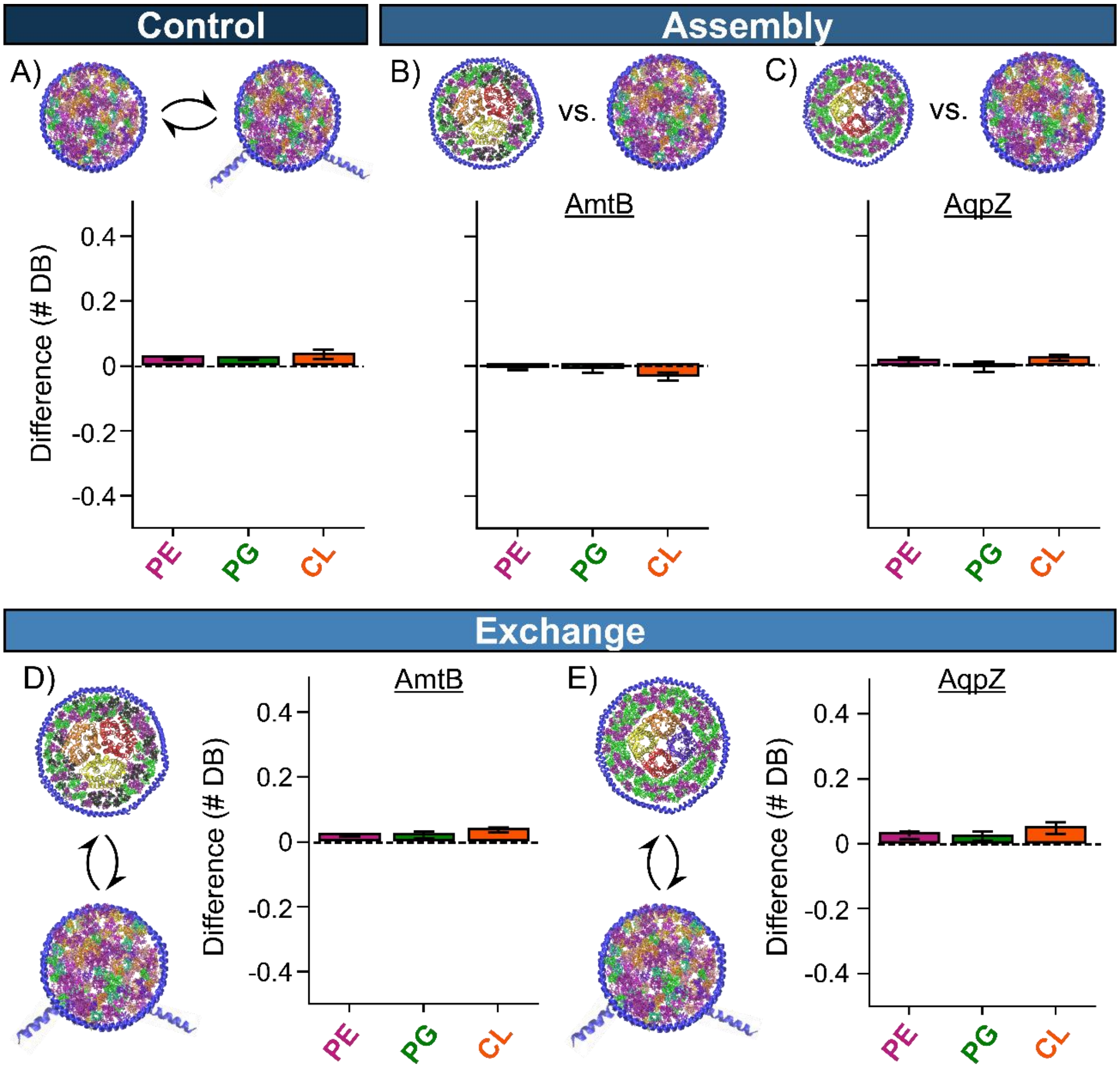

**Figure S3**. Changes in the *E. coli* lipid class average lipid tail unsaturation for each class weighted by the square root of their intensity for: (A) empty untagged versus empty tagged nanodiscs without membrane proteins after lipid exchange; nanodiscs assembled with AmtB (B) or AqpZ (C) versus empty untagged nanodiscs prior to lipid exchange; and AmtB (D) or AqpZ (E) untagged nanodiscs versus empty tagged nanodiscs after lipid exchange. Full details are provided in **Table S2**. Bar heights are the average difference weighted average tail unsaturation in total number of double bonds (# DB) for each class for three replicates with error bars showing standard deviations. Positive changes represent more double bonds in the empty untagged nanodisc population in A and membrane protein nanodisc population in B–E.

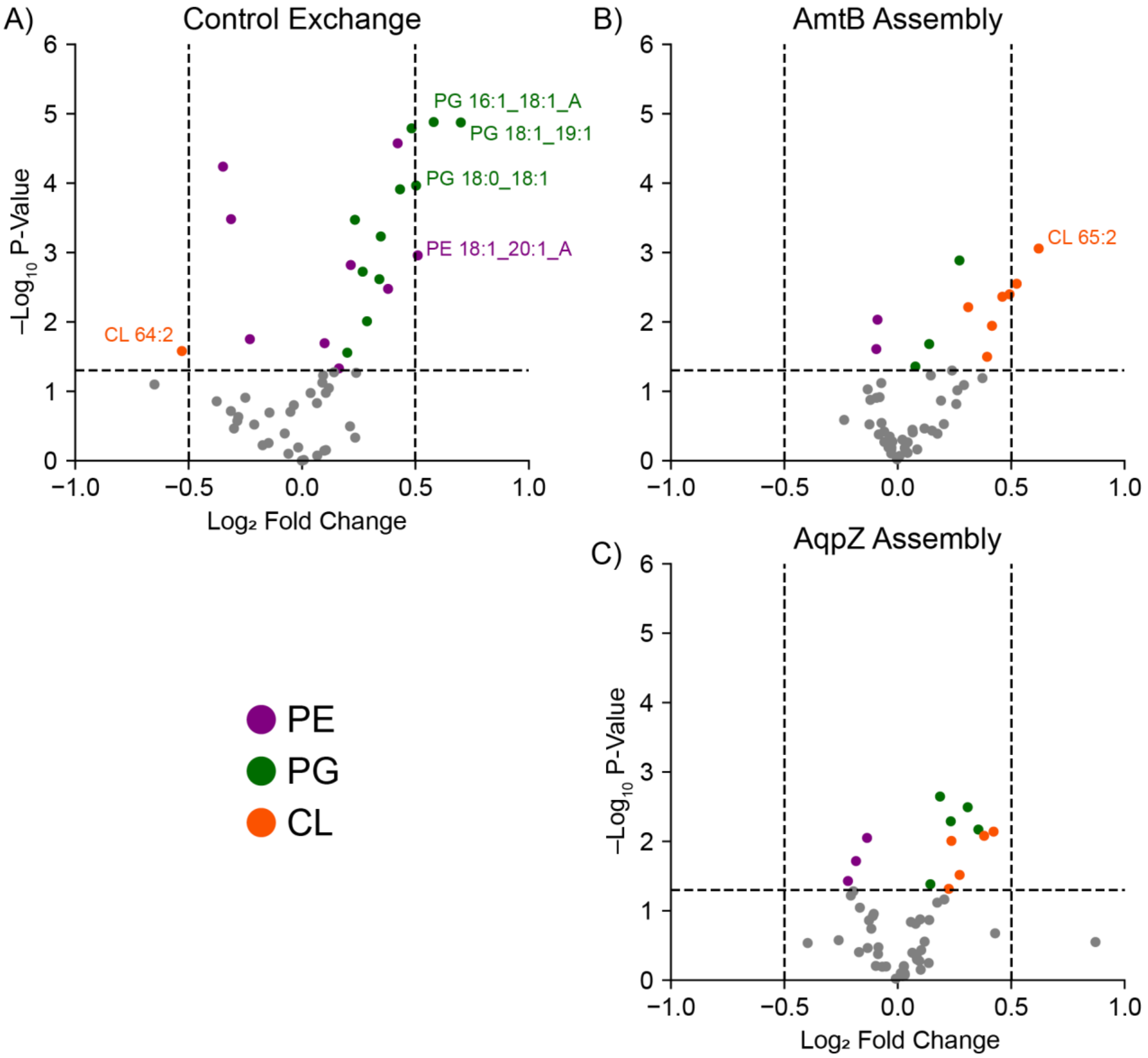

**Figure S4.** Individual changes in the identified *E. coli* lipids after lipid exchange for control nanodiscs without membrane proteins (A) and during nanodisc assembly prior to lipid exchange for AmtB (B) and AqpZ (C). Select lipids with the greatest fold changes and p-values are annotated. Full details are provided in **Table S4**. Positive $\log_2$ fold change values indicate higher levels in either empty untagged nanodiscs (A) or the membrane protein nanodiscs(B and C).

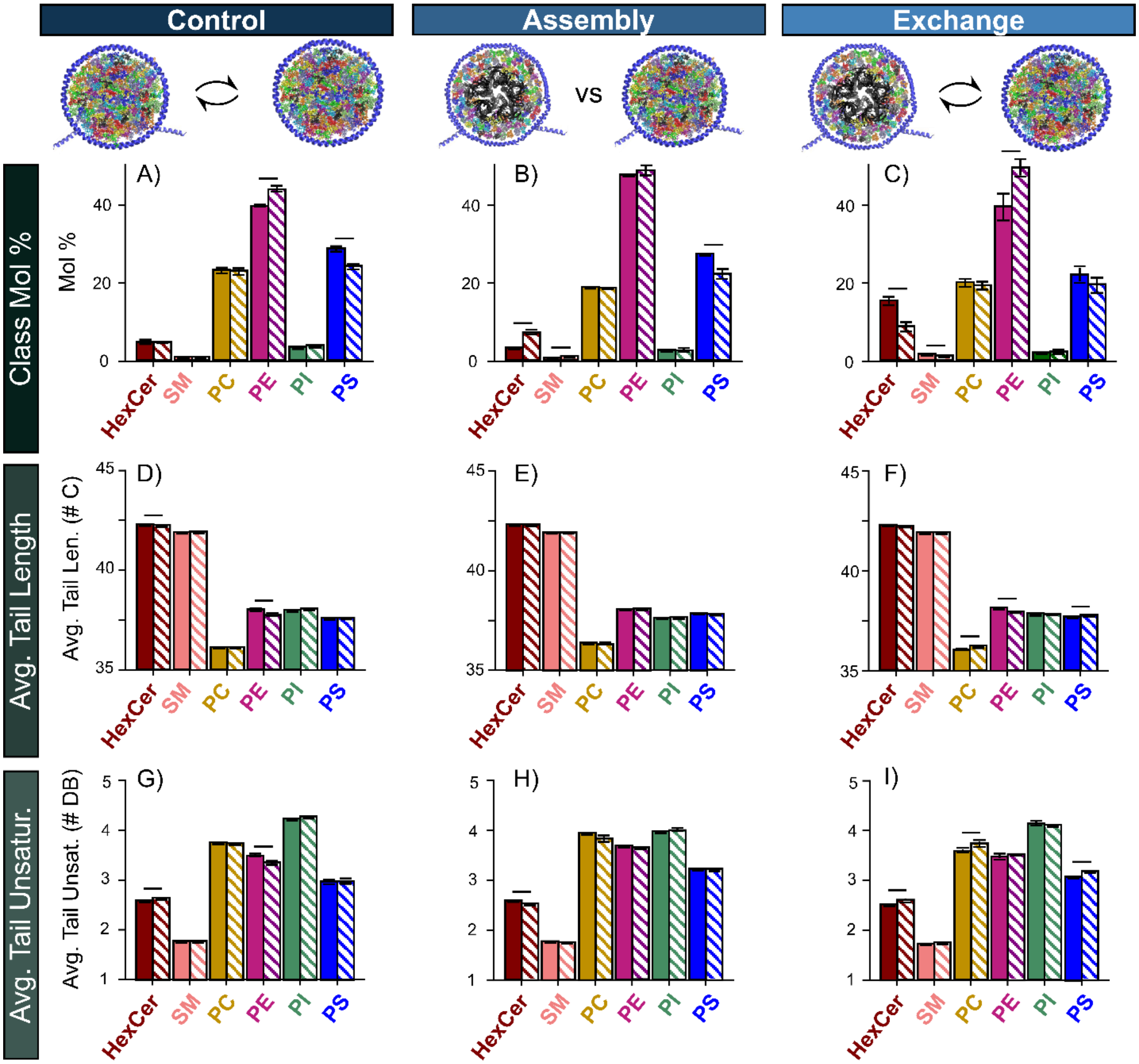

**Figure S5.** Absolute values for brain lipid extract relative class mole percent (A–C), the average total number of carbons in the fatty acid tails weighted by the square root intensity (D–F), and the total number of double bonds in the fatty acid tails weighted by the square root intensity (G–I) for: control lipid exchange without AChR (A, D, G), nanodisc assembly (B, E, H), and lipid exchange with AChR (C, F, I). Full results are provided in **Table S3**. Solid bars represent empty tagged nanodiscs in A, D, and G and empty untagged nanodiscs in all others. Shaded bars represent empty tagged nanodiscs in A, D, and G and AChR nanodiscs in all other panels. Significant ($p<0.05$) differences that exceed a 5% fold change threshold (A–C) or 0.05 difference (D–I) are marked with a horizontal line.

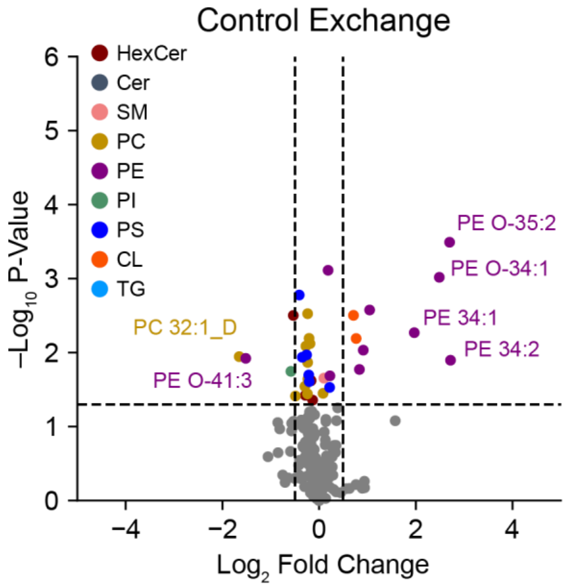

**Figure S6.** Individual changes in the identified brain lipids after lipid exchange for control nanodiscs without membrane proteins. Select lipids with the greatest fold changes and p-values are annotated. Full details are provided in **Table S5**. Positive $\log_2$ fold change values indicate higher levels in the empty tagged nanodiscs.

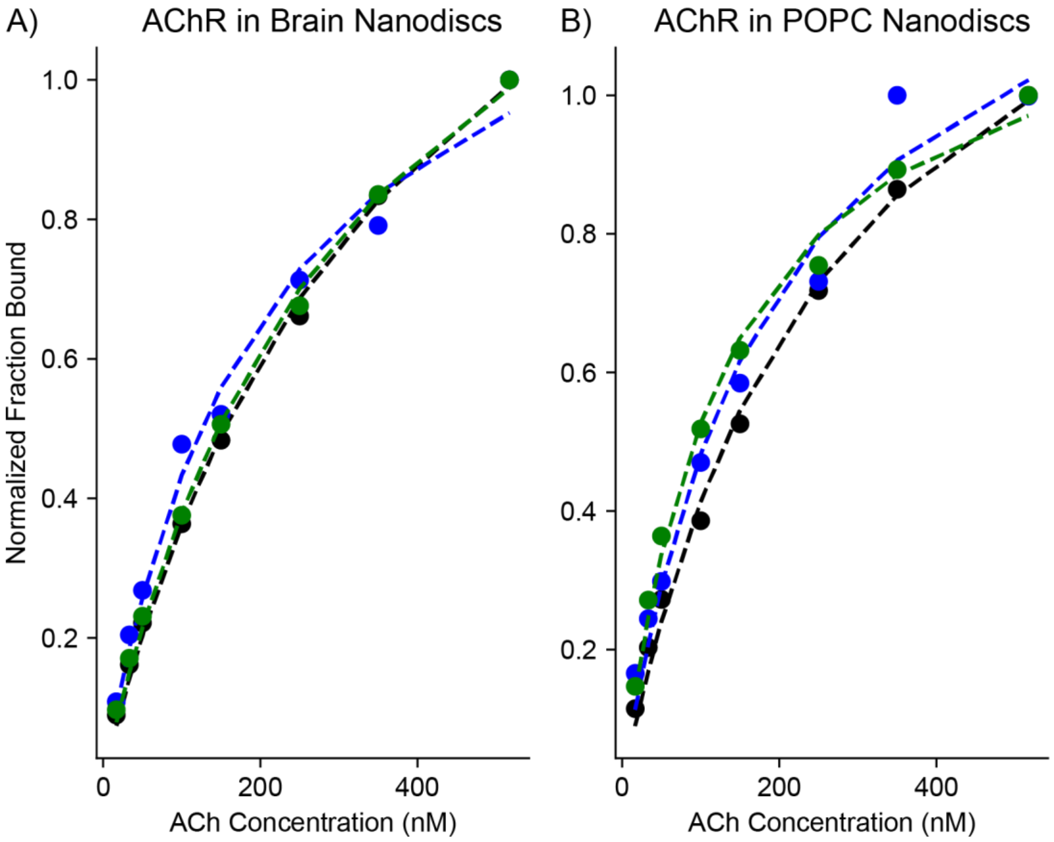

**Figure S7.** Fitting of nanoSPA$^{TM}$ for acetylcholine (ACh) binding in different nanodiscs. The scintillation response of the empty control nanodiscs was subtracted from the AChR in either brain lipid extract (A) or POPC nanodiscs (B), and they were normalized to the highest response. $K_d$ values were determined from the fit to be 299 ± 67 nM and 194 ± 53 nM, respectively.

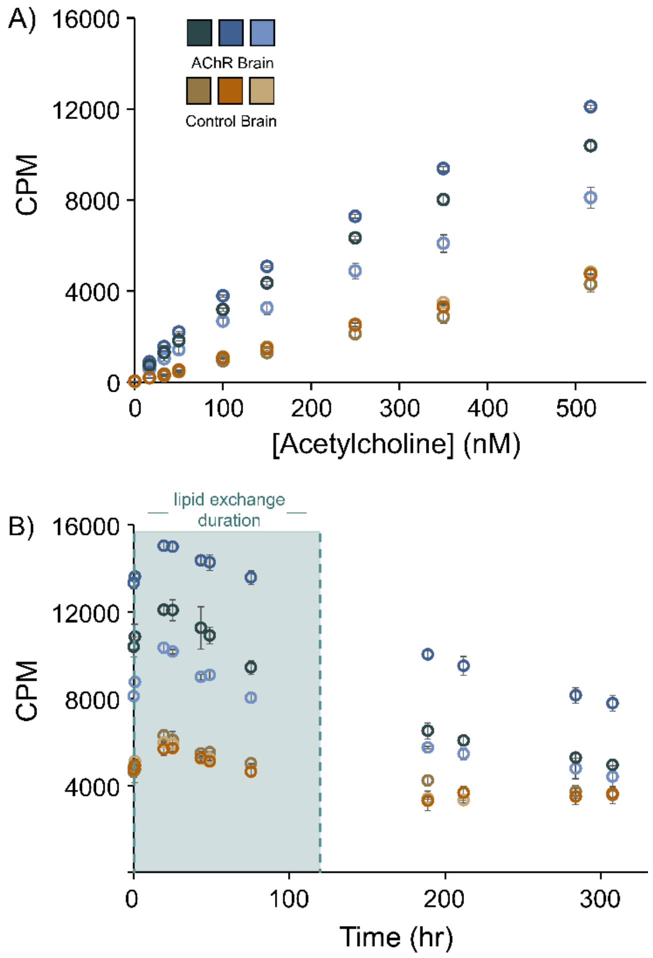

**Figure S8.** Scintillation proximity assay showing the counts per min (CPM) for three replicates of AChR in brain nanodiscs (blue) vs control brain nanodiscs without AChR (brown) as a function of $^{3}$H-acetylcholine (A) and at a fixed concentration over 12 days (B), which demonstrates the stability of AChR over the lipid exchange period.

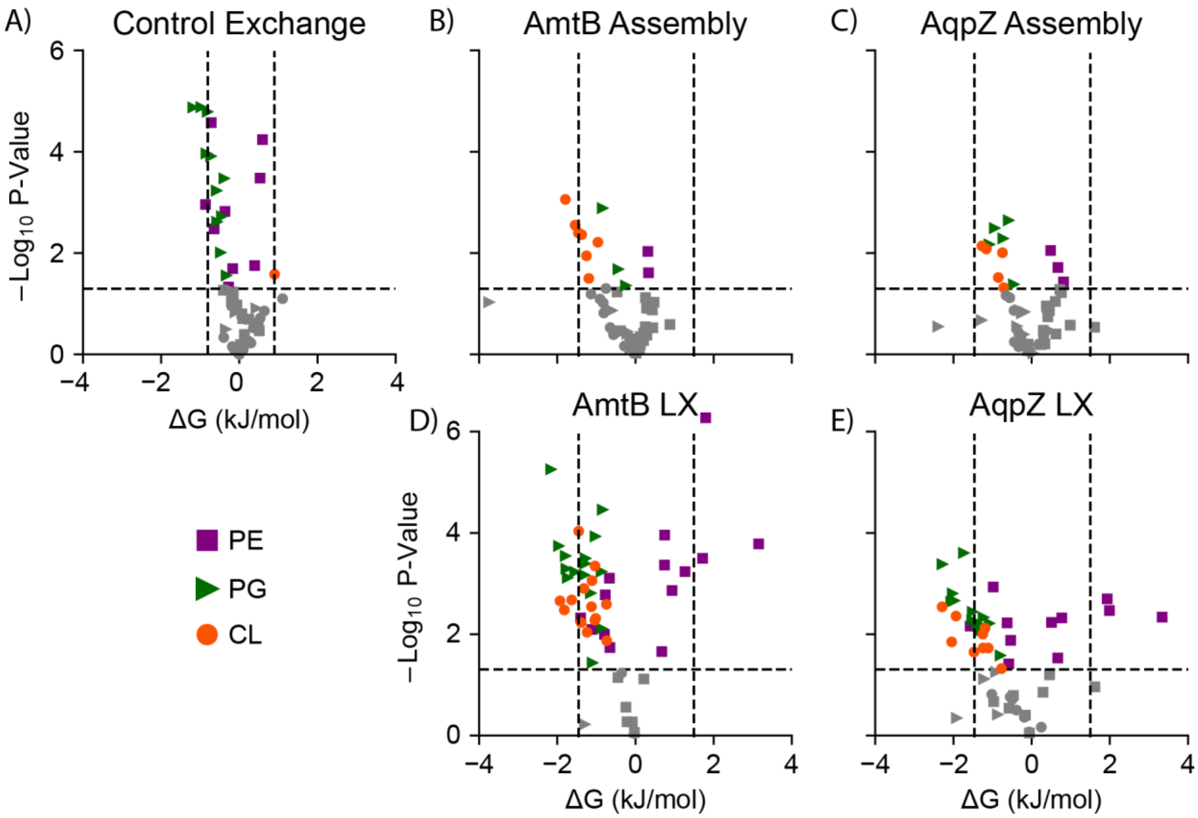

**Figure S9.** Thermodynamic analysis of individual lipid changes during the control lipid exchange with empty nanodiscs (A), AmtB assembly (B), AqpZ assembly (C), AmtB after lipid exchange (D), and AqpZ after lipid exchange (E). For annotation of select species, see **Figure 3**. Vertical dashed lines represent propagation of the 0.5 $\log_2$ fold change cutoff into ΔG space. Results are provided in **Table S6**.

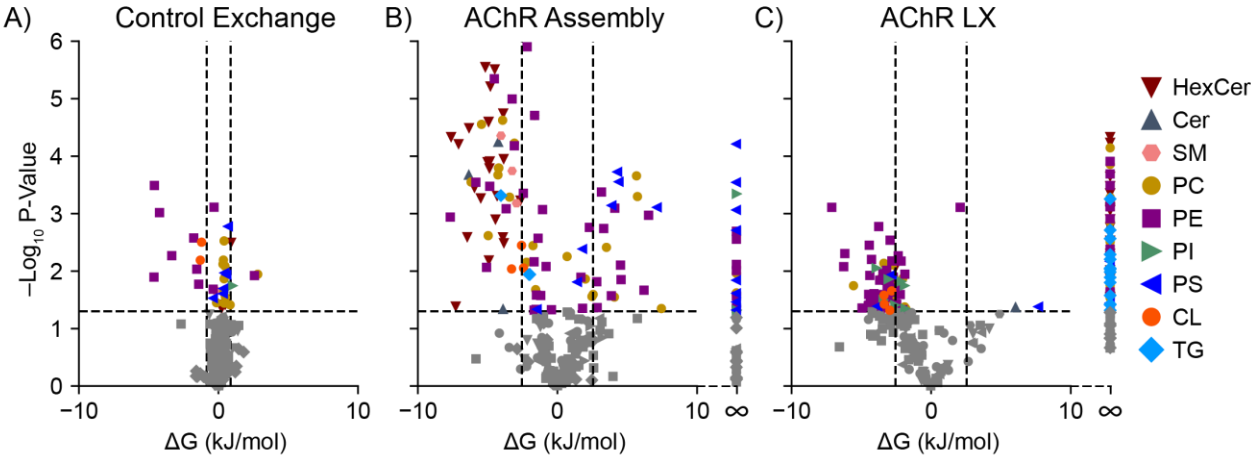

**Figure S10.** Thermodynamic analysis of individual lipid changes during the control lipid exchange with empty nanodiscs (A), AChR assembly (B), and AChR after lipid exchange (C). For annotation of select species, see **Figure 5**. Vertical dashed lines represent propagation of the 0.5 $\log_2$ fold change cutoff into ΔG space. Results are provided in **Table S7**.

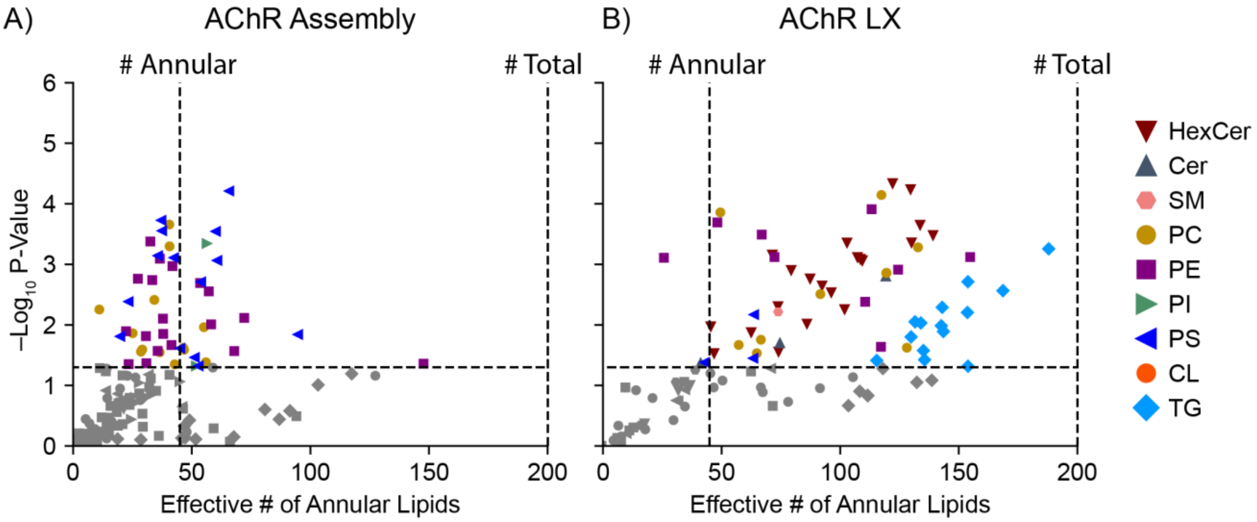

**Figure S11.** The effective number of annular lipids during AChR nanodisc assembly (A) and AChR after lipid exchange (B) calculated for all individual lipids with K < 1 using *Eq. S10*. For annotation of select species, see **Figure 5**. Vertical dashed lines represent the number of annular lipids previously measured (45)[14] and the total number of lipids predicted (200).
